# Silver nanoparticle induced oxidative stress augments anticancer gut bacterial metabolites production

**DOI:** 10.1101/658609

**Authors:** Prerna Bhalla, Raghunathan Rengaswamy, Devarajan Karunagaran, G. K. Suraishkumar, Swagatika Sahoo

**Author notes:** To whom correspondence should be addressed: G. K. Suraishkumar : Department of Biotechnology, Bhupat and Jyoti Mehta School of Biosciences building, Indian Institute of Technology Madras, Chennai 600036 INDIA;, and Swagatika Sahoo: Department of Chemical Engineering, and Initiative for Biological Systems Engineering, Indian Institute of Technology Madras, Chennai 600036 INDIA.

## Abstract

Colorectal cancer (CRC) is the fourth leading cause of mortality, world-wide. Gut bacterial dysbiosis being one of the major causes of CRC onset. Gut microbiota produced metabolites, e.g. folate and butyrate play crucial roles in cancer progression and treatment, and thus, need to be considered for effective CRC management. A potential cancer therapy, i.e., use of silver nanoparticles (AgNPs), imparts cytotoxic effects by inducing high intracellular reactive oxygen species (ROS) levels. However, the simultaneous interactions of AgNPs with gut microbiota to aid CRC treatment has not been reported thus far. Therefore, in this study, variation of intracellular ROS concentrations, in *Enterococcus durans* (*E. durans*), a representative gut microbe, was studied in the presence of low AgNP concentrations (25 ppm). Increases were observed in intracellular hydroxyl radical and extracellular folic acid concentrations by 48% and 52%, respectively, at the 9^th^hour of microbial growth. To gain a systems level understanding of ROS metabolism in E. *durans,* genome scale metabolic network reconstruction and modeling was adopted. *In silico* modeling reconfirmed the critical association between ROS and folate metabolism. Further, amino acids, energy metabolites, nucleotides, and butyrate were found to be important key players. Consequently, the anticancer effect of folic acid was experimentally studied on HCT116 (i.e., colon cancer cell line), wherein, its viability was reduced to 79% via folate present in the supernatants from AgNP treated *E. durans* cultures. Thus, we suggest that the inter-relationship between gut bacteria and AgNP-based cancer treatment can be used to design robust and effective cancer therapies.

## Introduction

Colorectal cancer (CRC)^1^ is characterized as a chronic consequence of abnormal gut bacterial metabolism and constitution (1). It is the fourth leading cause of cancer related mortality (2), and is defined as a complex association of tumor cells, non-neoplastic cells and a large amount of gut microbes such as *Streptococcus bovis*, *Helicobacter pylori*, *Enterecoccus faecalis etc.* that render carcinogenic effects (1, 3, 4).Various factors such as infection, diet and lifestyle can disrupt the symbiotic association between the host and the resident gut microbiota, leading to cancer (4).

The gut microbiome is composed of microbes that inhabit the gastrointestinal tract (GIT), which aid in human digestion, and serve as a metabolically active organ (5, 6). *Bifidobacterium* and *Bacteriodes* are the most abundant gut microbial species inhabiting the human gut (6). Gut microbes are also markers of homeostatic alterations often associated with chronic diseases. Perturbations in the composition and metabolic activity of the gut microbes, which are facultative or strict anaerobes, often result in neurodegenerative disorders (7) and cancer (4, 8).

The gut microbiome that consists of facultative or strict anaerobes (4, 8, 9) synthesizes and supplies essential nutrients such as short chain fatty acids (SCFAs) including butyrate, vitamin K and B group vitamins (B12, folate, riboflavin) (10). Amongst these metabolites, SCFAs and folic acid have been reported to be of importance in CRC initiation, progression and treatment (11, 12). Butyrate acts as a carbon source for colonocyte growth/maintenance (13), and protects against colon carcinogenesis, by modulating cancer cell apoptosis, cell proliferation and differentiation by inhibiting histone deacetylase (14). On the other hand, folic acid maintains the genetic integrity of normal colonocytes, and plays a crucial role in CRC management and treatment (it is often used as an adjunct with chemotherapeutic drugs to suppress tumor growth) (15).However, the effect of low concentrations of folic acid on cancer cell viability has not yet been reported. This study reports the unexpected effect of low folic acid levels on HCT 116, a colon cancer cell line.

Conventional cancer therapies such as chemotherapy and radiotherapy induce cytotoxic effects on the healthy cells, as well, due to lack of selectivity or high dosages (16). Nano-biomedicine is emerging as a novel anti-tumor therapeutic alternative (17), which can reduce the above side-effects on healthy cells. But nanoparticles at high concentrations can be toxic to human cells, predominantly due to lethal oxidative stress (18). However, at low concentrations (e.g. 32 ppm) silver nanoparticles (AgNPs), showed no significant toxicity to normal cells (19), and thus, can be of immense therapeutic importance. The impact of the interactions between gut microbes and AgNPs to aid CRC treatment has not been explored so far. Nanoparticles (NPs) induce cytotoxicity in the cancer cells by generation of oxidative stress due to excessive reactive species (RS) (18, 19), or by disrupting the cell membranes (20). RS are a class of highly reactive chemical species consisting of reactive oxygen species (ROS), e.g. superoxide, (O_2_^−^), hydroxyl radicals (OH), reactive nitrogen species (RNS), and others (21). Induced ROS, at appropriate levels, can increase metabolite production (31, 32), and hence, they can be expected to improve the production of anti-cancer metabolites (such as folate, butyrate) from gut bacteria. Thus, AgNPs, when administered for colon cancer treatment, are expected to be effective by two mechanisms: (i) by directly inducing cytotoxicity in cancer cells, and (ii) by increasing the anti-cancer metabolite production by gut bacteria through induced ROS (Fig. 1). The role of gut microbiota as a producer of folic acid or other anti-cancer metabolites, in the presence of oxidative stress has not been studied. Among the members of the *Enterococci* genus, *E. durans* is classified as a non-pathogenic inhabitant of the colon, rarely associated with human infections (22). Recent studies (23) suggest that *E. durans* is a beneficial gut resident because it is a potential anti-inflammatory probiotic in gut inflammatory disorders therapy. This study reports the effects of AgNP generated oxidative stress on metabolism of *E*. *durans*, which were investigated through experiments and further refined and directed via genome-scale metabolic modeling approach.

**Figure 1:**
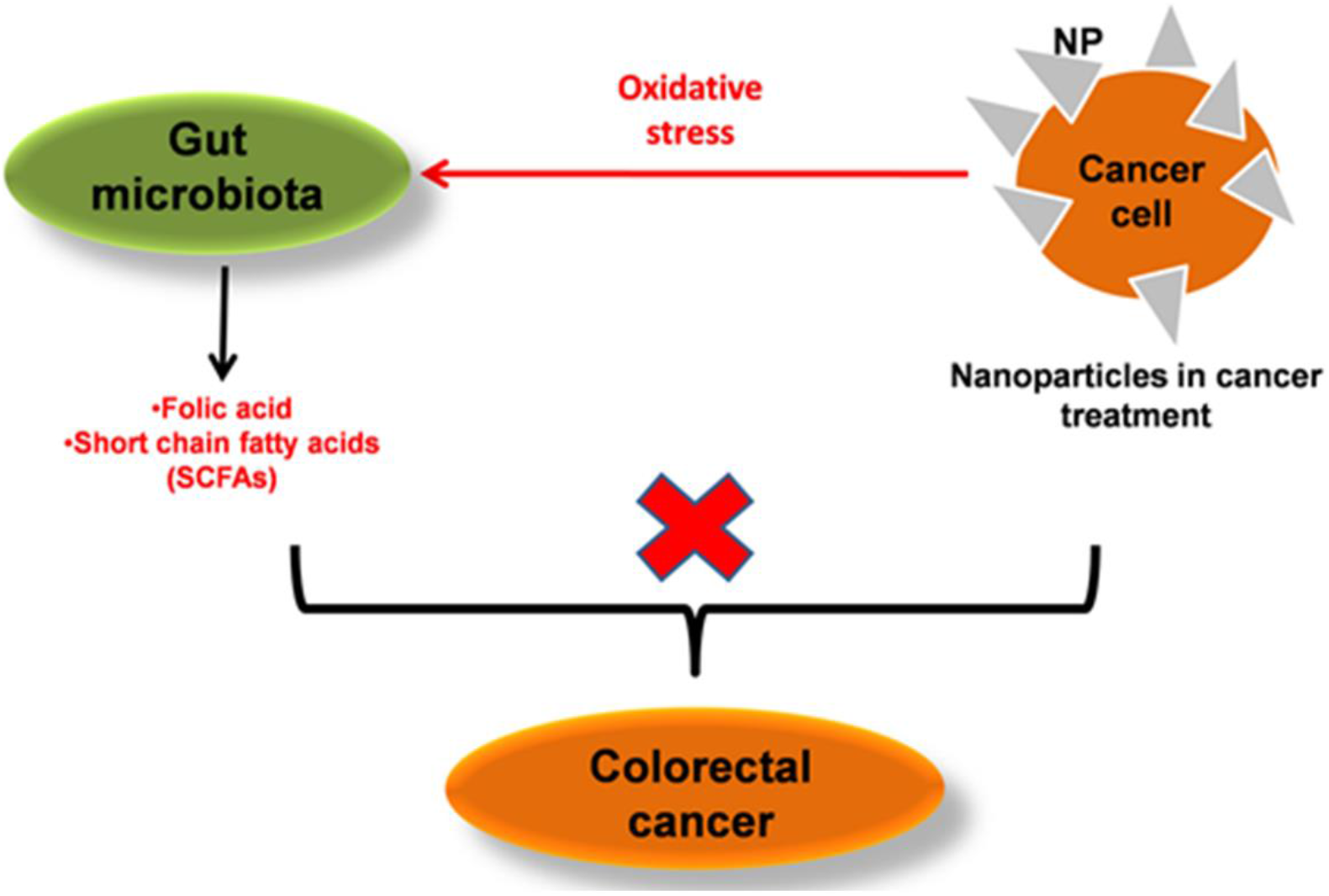
An overview of the direct and indirect effects of AgNPs mediated oxidative stress on gut bacterial metabolism in treatment and management of CRC. AgNPs are known to mediate their direct cytotoxic effects through increased generation of reactive species inside the target cells. Besides, the AgNP-generated oxidative stress can also affect the cellular metabolism and certain pathways of interest (in this case, the anti-cancer metabolites) thereby enhancing indirect cytotoxicity.

Constraint-based reconstruction and analysis (COBRA) method remains the most preferred systems biology tool to interrogate phenotypic properties of the target organism (37). Genome-scale metabolic network (GEMs) integrates the biochemistry, genomic, and genetics of the modeled organism, and serves as the best platform to gain a systems understanding of the cellular /molecular phenomenon (24). GEMs for numerous human microbes have been compiled successfully using COBRA framework (25). The effect of changed dietary components and environments on cellular robustness can be readily simulated and analyzed (26). Recently, COBRA models of 773 most prominent gut microbes were made available through AGORA (Assembly of gut organisms through reconstruction and analysis). AGORA is a resource of GEMs that is effective to understand metabolic diversities of microbial communities (27). These large scale metabolic networks, so far, have not taken into account the probable effects of reactive species on the gut bacterial metabolism.

In this study, the effects of AgNP induced oxidative stress on the production of anti-cancer metabolites (i.e., folic acid and butyric acid) was investigated in *Enterococcus durans*. A combinatorial extensive wet lab experimentation and genome-scale metabolic modeling approach was adopted to analyze the (i) impact of ROS on the cellular metabolism of the bacterium, and (ii) the cytotoxic potential of low folic acid levels as potential cancer treatment strategy.

## Results

### 2.1 Characterization of AgNPs and their interaction with E. durans

The sizes and stability of AgNPs in solution were first characterized using dynamic light scattering (DLS) and zeta potential measurements, respectively. The average particle size was 185±6 nm, and the zeta potential of the bacterial medium with nanoparticles was −13±0.95mV. The topographic details and composition of nanoparticle treated bacterial cells were examined through scanning electron microscopy (SEM). The interaction between nanoparticles with the bacterial cells was observed in the SEM micrographs (Fig. 2). The composition analysis showed 9.7% Ag content in the biomass.

**Figure 2:**
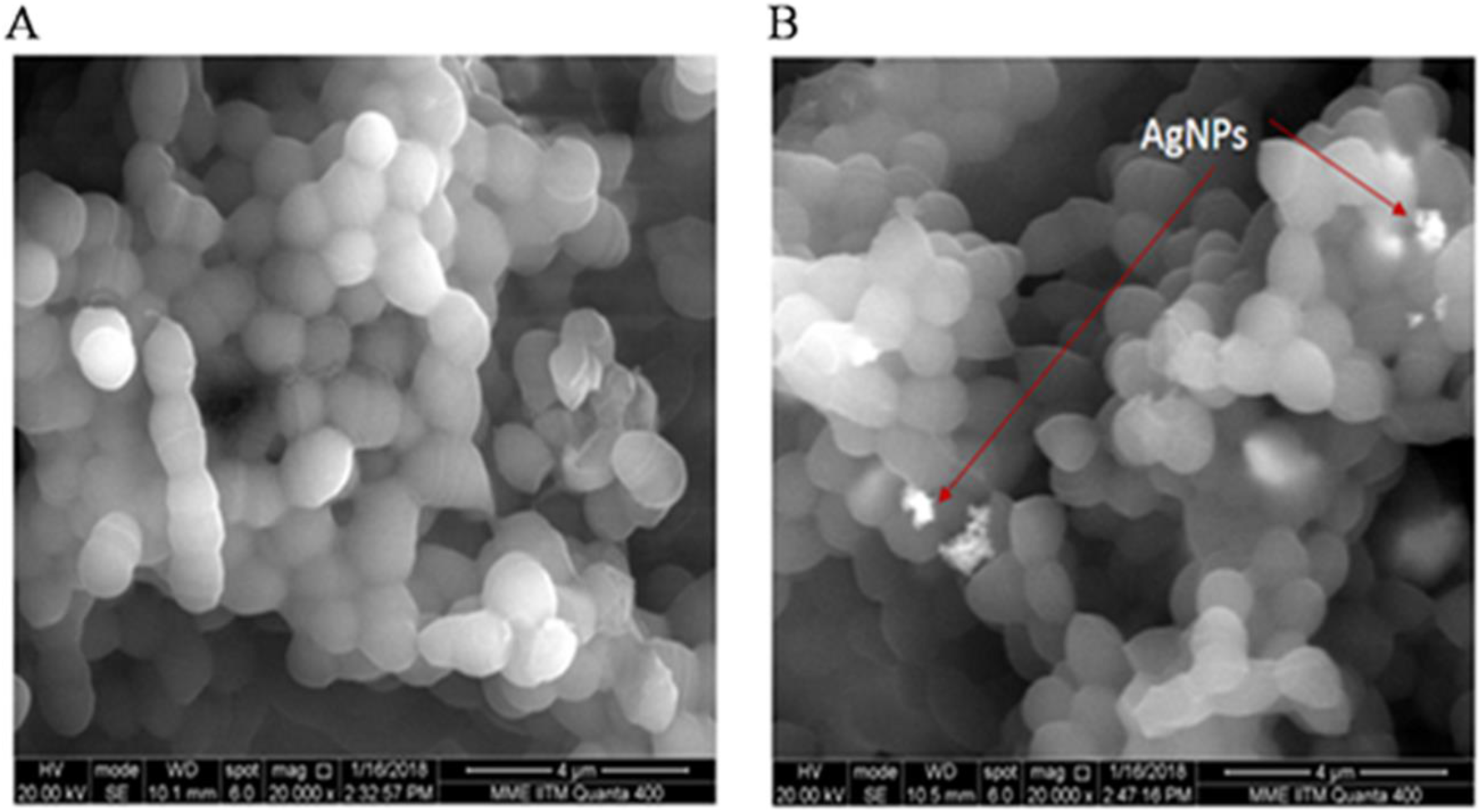
Scanning electron microscopy to elucidate the interaction between silver nanoparticles (AgNPs) and bacterial cells. 2(A) represents bacteria alone, whereas, 2(B) shows AgNPs interacting with the bacterial cells

### 2.2 Lower concentrations of silver nanoparticles had no major deleterious effects on E. durans

The above characterizations provided the information on the relevant properties of AgNP. The effect of AgNP concentration on the growth and cell viability of *E. durans* was next studied by exposing the bacterial cultures to different concentrations of AgNPs between 25ppm to 250ppm (Fig. 3). The specific growth rate for the cultures treated with lowest concentration (25ppm) of nanoparticles was 0.198±0.03 h^−1^. The specific growth rate reduced by 8% at 25 ppm compared to control. Further, the specific growth rate decreased with increasing concentration of nanoparticles. At 250 ppm concentration, the specific growth rate was 0.08±0.016 h^−1^, which indicates the deleterious effects of nanoparticles at high concentrations. However, the lower concentrations of AgNPs in the range studied did not adversely affect the growth of the organism.

**Figure 3:**
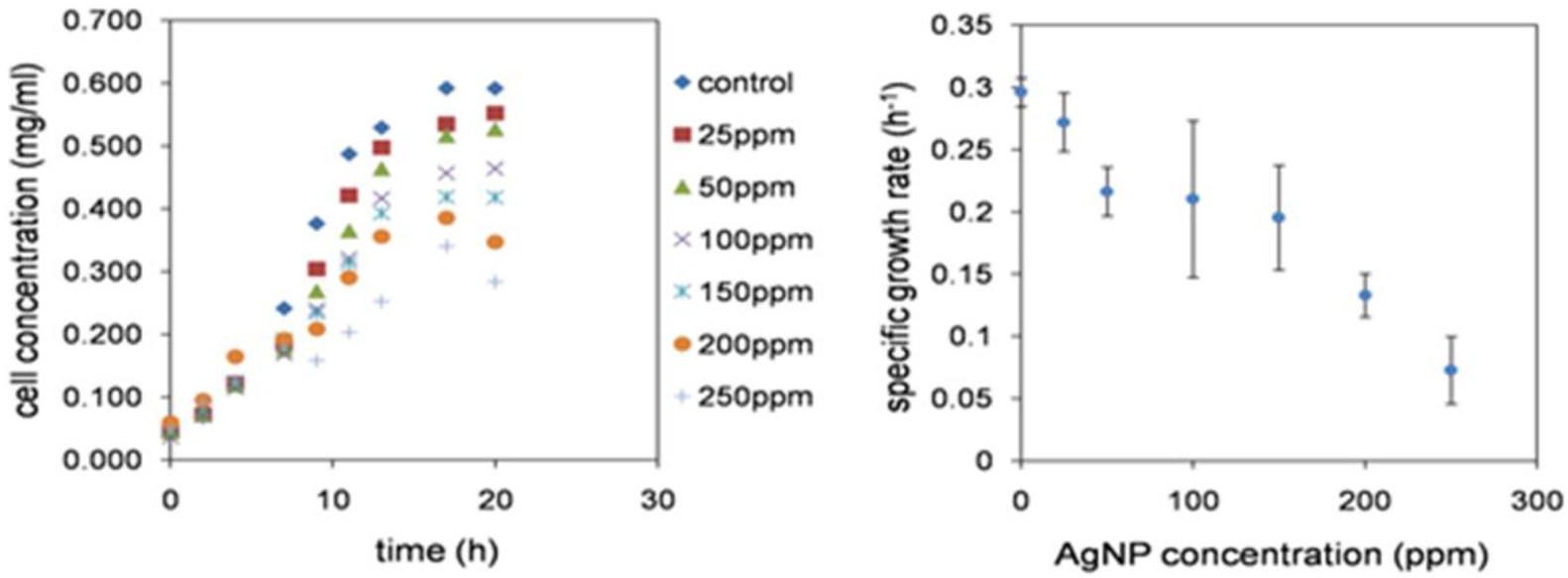
Variation of E.durans growth with different concentrations of AgNPs. Exposure of bacterial cultures to higher concentrations of the nanoparticles results in excessive cell death. This is confirmed by significant decrease in the specific growth rates of bacterial culture with increasing AgNP concentration. The specific growth rate, is a function of cell viability (maximum growth) with respect to time, is also affected negatively at higher concentrations of AgNPs.

### 2.3 Exposure of E. durans to lower AgNP concentration increases intracellular ROS generation

Although the AgNPs at 25 ppm did not significantly affect growth, they may alter the relevant aspects of cellular metabolism. For example, nanoparticles are known to induce intracellular ROS generation, viz. superoxide and hydroxyl radicals, inside bacterial cells (28).The intracellular levels of superoxide and hydroxyl radicals were quantified at 25ppm, and the results are presented in Fig. 4. At the 6^th^hour (mid log phase) of nanoparticle treated bacterial culture growth, 0.273±0.01 nanomoles/(g-cell) intracellular superoxide concentration was generated, thereby showing an increase by 13% in the specific superoxide level, when compared with control. The specific intracellular hydroxyl radical level showed48% increase (1.057±0.02 nanomoles/g-cell) in the late-log phase, when compared to control. Thus, intracellular reactive species levels were altered at lower AgNP concentration, without affecting the cell viability.

**Figure 4:**
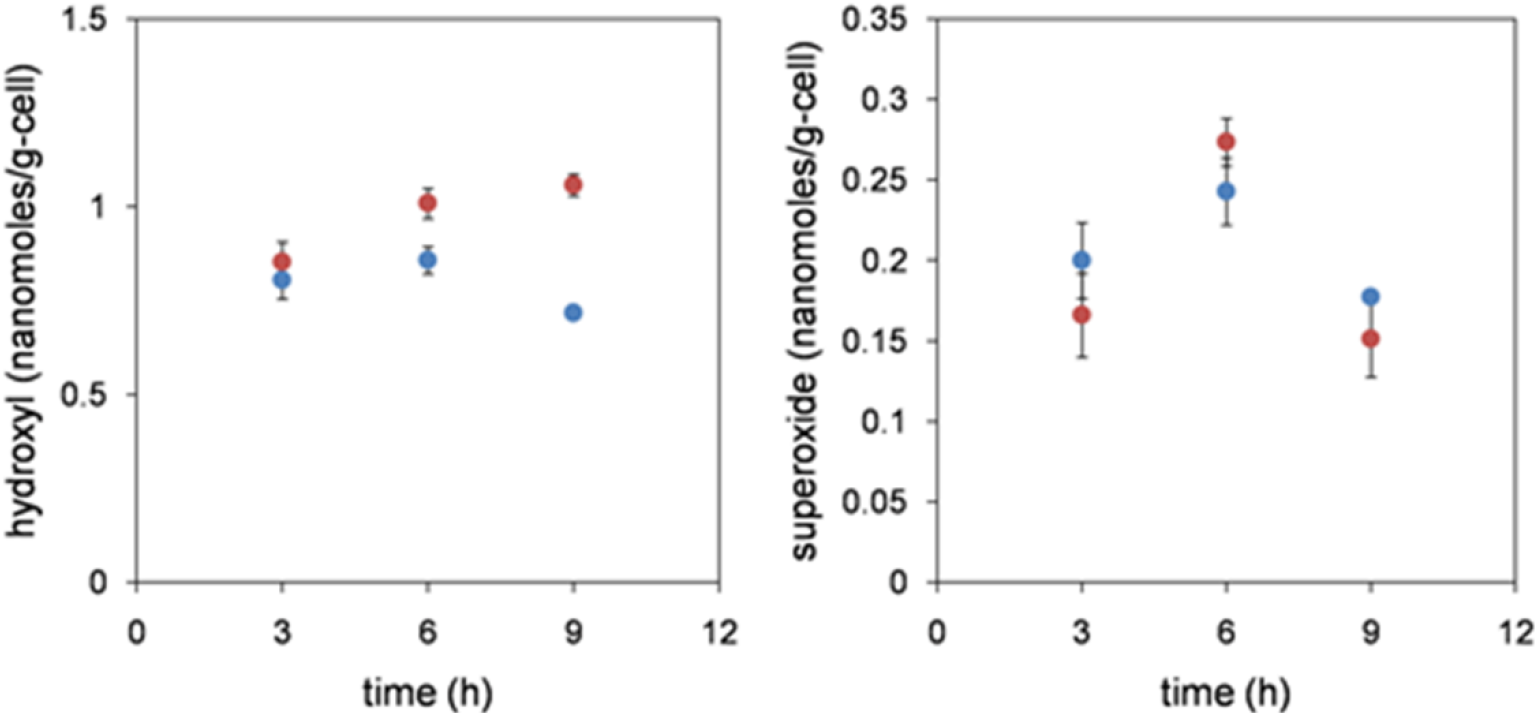
Reactive species time profile in the absence and presence of AgNP generated oxidative stress (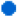 indicates control culture; 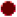 indicates culture treated with 25 ppm AgNP).The hydroxyl radical concentration in the presence of lower concentration of AgNP, showed increase with increase in time of exposure, when compared to the control. The superoxide radical levels, increased during the mid-log phase of growth (on exposure to AgNPs), and then dropped, when compared to control.

### 2.4 Reactive oxygen species generation alters intra- and extra-cellular folate concentrations in bacterial cells

We then studied the effects of nanoparticle (at 25 ppm) mediated, increased oxidative stress on bacterial folic acid production (Fig. 5). Three different time points, one each in the lag (3^rd^ h), mid-log (6^th^ h), and late-log (9^th^ h) phases were chosen to measure the intracellular and extracellular concentrations of folic acid using HPLC. It was also observed earlier that specific extracellular folic acid levels reduced after the 9^th^ hour, and hence, only the above points were chosen. The maximum difference in the intracellular folic acid levels produced by treated cells and control was observed at the 6^th^ hour of microbial growth. The intracellular folic acid level was 45.53±0.012 nanomoles/g-cell in nanoparticle treated culture, which was 49% higher compared to control.

**Figure 5:**
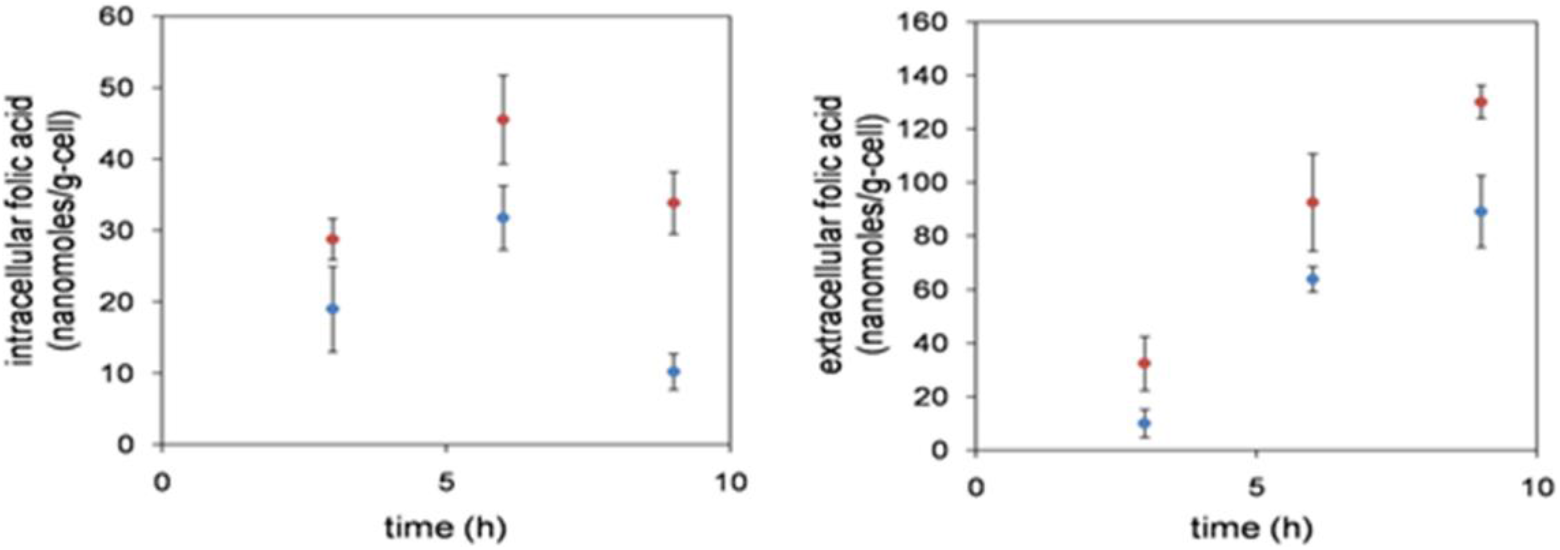
The intracellular and extracellular folic acid concentrations increased on exposure to AgNPs (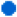 indicates control culture; 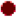 indicates culture treated with 25 ppm AgNP). This increase could be an outcome of the oxidative stress induced in the organism, which in turn impact folate related metabolic pathways. However, higher concentrations of AgNPs negatively affected folic acid generation by causing cell death.

Moreover, *E. durans* was found to secrete folic acid into the extracellular space. The maximum specific level of extracellular folic acid was detected at the 9^th^ hour in nanoparticle treated cultures (128.84±0.16 nanomoles/g-cell), which was 52% higher compared to control.

### 2.5 In silico analysis of reactive species provides insight into the alteration of gut microbial metabolism

Genome-scale metabolic modeling of *E. durans* was used to identify the important metabolic consequences associated with increased generation of reactive species (ROS/RNS), within the microbial system. To assess the fundamental effects of nanoparticle generated reactive species on folic acid metabolism in *E. durans,* a constraint-based metabolic modeling approach was adopted. The genome-scale metabolic model of *E. durans* was obtained from Virtual Human Metabolic (VMH) database (27) and further expanded with reactive species reactions (Table 1, Supplement Table S1). The relevant reactive species reactions were obtained through literature search. These reactions focused on the bio-chemical interactions between reactive species (predominantly superoxide, hydroxyl radical and nitric oxide) and major bio-molecules such as amino acids and nucleic acids. The input constraints imposed on the model were based on the experimental conditions provided for microbial growth. For instance, the model was studied under aerobic conditions (*E. durans* being a facultative anaerobe), with glucose as a major carbon source present in the chemically defined media used to culture the bacteria.

**Table 1:**
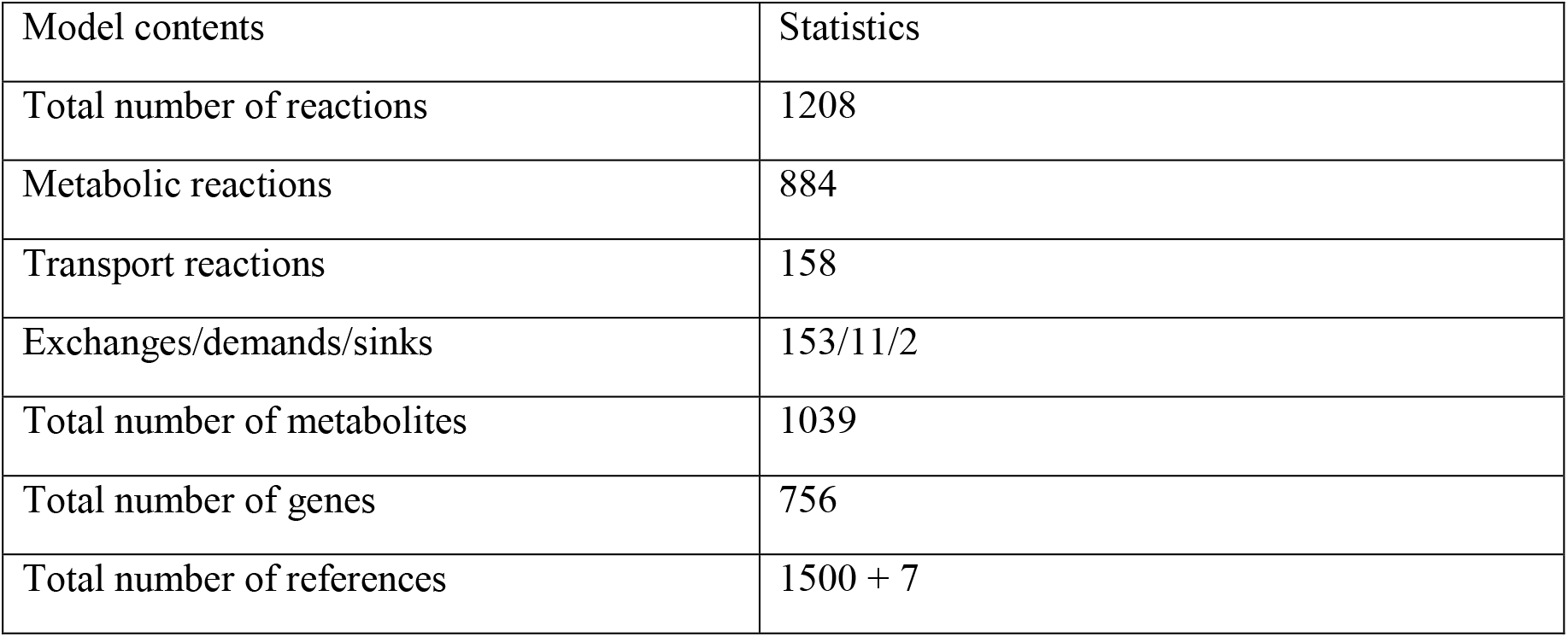
Statistics of genome scale metabolic model of E. durans containing ROS reactions and related metabolites.

Further, to investigate the effects of ROS on metabolic network of the microbe, flux variability analysis (FVA) was performed. The relative changes in the network fluxes before and after the addition of ROS reactions were evaluated through the flux span ratio (FSr). The FSr corresponds to the variability of each network reaction in the presence and absence of oxidative stress, under the defined constraints. The reactions having FSr values in the range 0.8>FSr>2, were most affected in the presence of ROS.

A total of seven reactions were found to be in the specified FSr range. These reactions were explicitly associated with folate metabolism and showed positive fluxes in the presence of reactive species. MinNorm analysis indicated that these seven reactions in turn influenced different major metabolic pathways like, amino acid/peptide metabolism, nucleotide metabolism, carbohydrate metabolism (pyruvate metabolism, glycolysis/gluconeogenesis) and energy metabolism, as elucidated in Fig.6.

**Figure 6:**
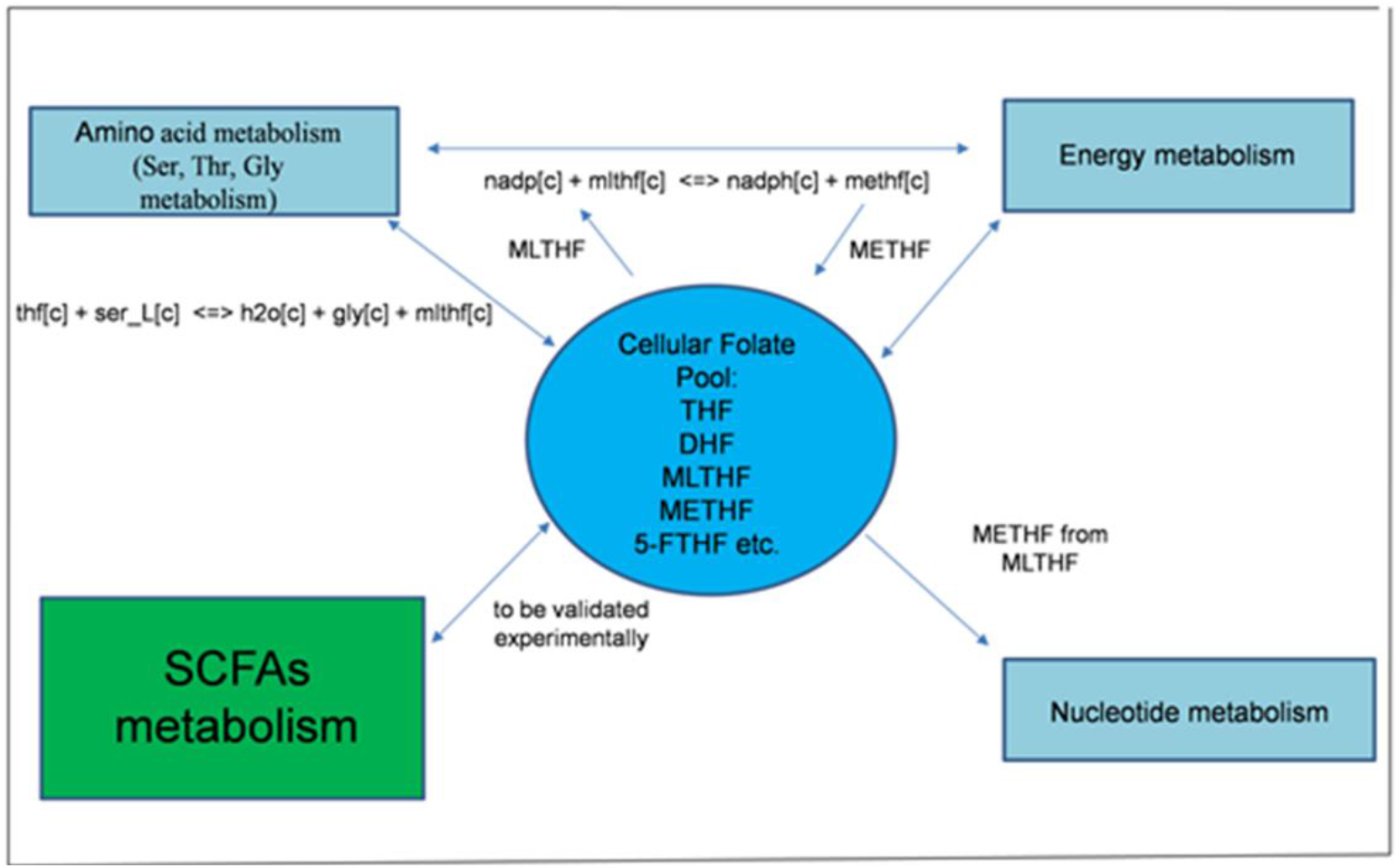
Folate metabolism and its association with different central metabolic pathways (viz. amino acid, energy and nucleotide metabolism), where the model predictions agreed with experimental evidences in literature. The link between SCFA metabolism and folic acid metabolic pathways is a novel model prediction, and needs to be experimentally validated.

### 2.6 Microbial model under oxidative stress exhibited secondary effects on butyrate-linked metabolic pathways

Although the associations between folic acid derivatives and energy metabolism, amino acid metabolism or nucleic acid metabolism are known (details mentioned in discussion section), the association between folic acid derivatives and SCFA metabolism (i.e., butyrate), as predicted by the constraint based modeling approach, is unknown. More specifically, an association between the cellular folate pool and butyrate (SCFA) metabolism (Fig. 6), under oxidative stress conditions was predicted by modeling analysis (Table 1).

### 2.7 Folic acid exhibited cytotoxic effects on HCT 116 colon cancer cell line at lower concentrations

Although it is known that folic acid enhances cancer viability (29), the effects of lower concentrations of folic acid on cancer cells have not been reported in literature. We tested different concentrations of folic acid on the viability of HCT 116 colon cancer cells (Fig. 7(A)). As expected, higher concentrations of folic acid (5 – 30μM) resulted in increased viability of cancer cells. On the contrary, unexpectedly, lower concentrations (1 – 0.1 μM) reduced cell viability. The optimal-cytotoxic concentration was around 0.5 μM folic acid, which resulted in decrease of viability by 14% when compared with the appropriate control. The reduction in cell viability at lower concentrations of folic acid was statistically significant. A P-value of 0.001458 was obtained on performing single factor ANOVA. Further, t-test (two-sample assuming equal variances) generated a P-value of 0.027004, statistically supported the finding.

**Figure 7:**
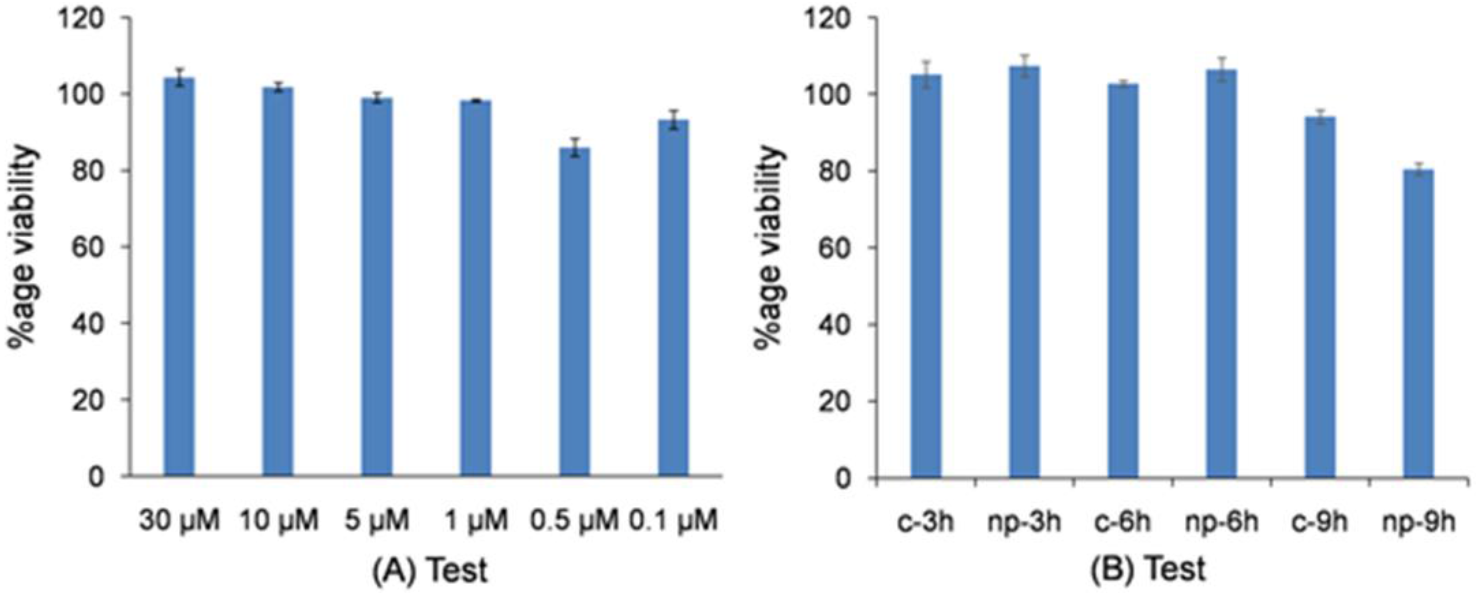
Effect of different folic acid concentrations on the viability of HCT116 cancer cells. (A) Lower concentrations of synthetic folic acid exhibited cytotoxic effects on cancer cells. (B) Also, MTT analysis of HCT116 treated with bacterial supernatants from cultures exposed to silver nanoparticles and control. Reduction in viability was observed for cells treated with 9th hour supernatants, the time point corresponding to maximum concentration.

### 2.8 Folate enriched supernatant from nanoparticles treated E. durans cultures affects cancer cells viability

Bacterial culture supernatant is rich in different metabolic secretion products. Many of these metabolites are known to have anticancer properties, some of which have been previously discussed. However, the role of folate/folic acid produced by gut microbiome in context of nanoparticles based targeting of colorectal cancer has not been studied yet. To observe the effects of secreted folic acid present in nanoparticle treated bacterial culture on HCT 116 cancer cell viability, the cancer cells were treated with crude bacterial supernatant. MTT assay was then performed. It was found that silver nanoparticles (25 ppm) treated bacterial culture supernatant exerted cytotoxic effects on cancer cells. The cancer cell viability treated with 9^th^ hour supernatant from nanoparticle treated cultures reduced to 79% of control (Fig. 7(B)), which is the time point corresponding to the release of the optimal concentration of folic acid produced in the culture. The reduction in cell viability on exposing the cancer cells to 9^th^ hour AgNP treated *E. durans* cultures was statistically significant. A P-value of 0.001429 was obtained on performing single factor ANOVA. Further, t-test (two-sample assuming equal variances) generated a P-value of 0.013265, statistically supported the finding.

## Discussion

Silver nanoparticles were characterized for their size and stability in the bacterial culture medium. The stability of the nanoparticles was assessed through zeta potential measurement. A negative value (–13±0.95 mV) indicated better stability of the dispersed nanoparticles. The higher stability also indicates that the well dispersed nanoparticles effectively, interacted with the bacterial cells in the culture, and affected their viability. Higher concentrations of AgNPs severely affected *E. durans* growth rate. This observation is in accordance with the literature studies, where it has been shown that AgNPs at higher concentrations induce their cytotoxic effects by disrupting bacterial cell membranes and permeability, thereby interfering with Na+ transport and homeostasis (20).The other significant explanation for the cytotoxic activity of AgNPs is the generation of reactive species, which results in inhibition of aerobic respiration in the organism and causes damage to its genetic moiety (18). However, AgNPs at concentrations as low as 25 ppm had minimal cytotoxic effects on *E. durans* viability. Since the context is use of nanoparticles for therapy, this lower concentration of AgNPs was chosen for all further experiments.

AgNPs have been reported to generate oxidative stress in a cellular system (37), hence, we studied the response of *E. durans* to nanoparticle treatment. The intracellular ROS (hydroxyl and superoxide radicals) levels were quantified at three different time points corresponding to different phases of bacterial growth. Despite *E. durans* exhibiting notable antioxidant properties (30), the lower concentration of AgNP resulted in increased generation of intracellular reactive species. The hydroxyl levels increased considerably during the late log phase in nanoparticle treated cultures, indicating that AgNPs are potent inducers of reactive oxygen species. The augmented reactive species levels are known to affect and damage the major bio-molecules, such as nucleic acids and proteins (18). This nanoparticle-generated oxidative stress, in turn, may also affect the cellular metabolic systems, thereby resulting in increased productivity of certain metabolites. For instance, previous studies in our laboratory have established the role of stress induced ROS in increased lipid accumulation in *Chlorella vulgaris* (31). Likewise, HOCl (a commonly used hydroxyl ion generator) treatment of *Xanthomonascampestris* improved the xantham gum productivity in different cultivations (32). In case of *E. durans,* the role of intracellular oxidative stress on the production of folic acid or other gut microbial metabolite has not yet been investigated. In our study, it was observed that the folate levels (both intracellular and extracellular) were modulated in response to the enhanced intracellular ROS levels, in nanoparticle (25 ppm) treated cultures. The intracellular folic acid level is a measure of the bacterial cell potential to produce and secrete extracellular folic acid, and thus needs to be quantified. The extracellular folic acid concentrations increased during the late log phase, which also corresponded to the time point of increased intracellular hydroxyl levels. This finding strongly supports the role of ROS in regulating the generation of microbial metabolites. At higher nanoparticle concentrations (> 50 ppm) the cell viability was reduced. Also, no detectable intracellular or extracellular folic acid levels were observed at AgNP concentrations above 50 ppm, which suggests a deleterious effect of high AgNP levels on folic acid synthesis and secretion.

The gut microbiota is a reservoir of many essential metabolites required by the host homeostasis (10). Amongst all these metabolites, folic acid (vitamin B group) is of major relevance due to its involvement in chief metabolic pathways (34, 35). To better understand the mechanism of the effects of reactive species on microbial metabolism, we used computational analysis. For this purpose, constraint-based metabolic modeling was used to analyze the systems level effects of ROS on the metabolism of *E. durans.* In our study, COBRA modeling helped establish the critical link between reactive species with other important intermediary metabolisms (i.e., amino acid, energy, and lipids) and their consequent effects on microbial metabolism. Previous studies involving systems biology approaches have helped in deciphering the host-microbe interactions and their consequences on human health. The reconstruction of genome scale metabolic networks under various constraints (mimicking diseased state), have successfully predicted the host phenotypic characteristics (25). A similar approach has been used to study the metabolism of gut microbe in presence of oxidative stress. As stated earlier, folic acid (and its derivatives like THF, DHF etc.) and SCFAs, such as acetate, propionate, valerate and predominantly butyrate, are of metabolic importance in the development or suppression of colon cancers. Additionally, butyrate is known for its tumor suppressive role in colon and stomach cancer (14).The increasing or diminishing levels of intracellular reactive species in bacteria, affect the generation of these metabolites of interest. So, it is crucial to understand how the entire metabolic network of the gut microbe gets affected in response to oxidative stress (Table 2). For instance, one of the reactions affected by the addition of ROS to the model system involved glycine, serine and threonine metabolism. In this reaction, the 3-carbon serine serves as one of the major sources in transferring one carbon moiety to tetrahydrofolate (THF) to form 5, 10-methylene THF (MLTHF) via glycine hydroxymethyl transferase (E.C.2.1.2.1) (34), A positive flux (500.5 mM/g-dry weight h) was generated through this reaction, thus implying an increased release of the folate derivative which contributed to the folate pool by affecting folate metabolism. In another case, NADPH and serine metabolism have also been associated with folate metabolism (35). MLTHF from serine metabolism is used to synthesize METHF (methenyl THF) via methylenetetrahydrofolate dehydrogenase (E.C.1.5.1.5), which in turn is utilized for nucleotide synthesis. The model with ROS reactions showed increased flux through this reaction, thus, supporting the involvement of folic acid in energy and nucleic acid metabolism, as well as, highlighting the role of reactive species in heightened folate production. Interestingly, the link between folic acid derivatives and butyric acid (SCFA) metabolism was revealed through the modeling predictions. Thus, this approach involving constraint based metabolic modeling served two purposes. Firstly, the curated model can facilitate studying of the effects of reactive species generated by AgNPs on the entire metabolic network of the gut microbe. Secondly, this methodology can be used to identify the metabolic pathways significantly associated with generation of anti-cancer metabolites in the presence of reactive species to improve upon the cancer treatment strategies. This is the first study of its kind that validates the gut bacterium metabolic model experimentally, where folate levels showed an increase in the presence of AgNP generated intracellular oxidative stress. The modeling results primarily emphasized the effects of reactive species on folate metabolism and generation, which may play a crucial role in strategizing colorectal cancer treatment therapies.

**Table 2:** Different folic acid reactions being affected on addition of ROS reactions to the unconstrained E.durans model that are validated in literature.

| S.No. | Reaction abbrev. | Flux<br>(mmol/gDW/hr) | Comments |
| --- | --- | --- | --- |
| 1. | glycine<br>hydroxymethyltransferase,<br>reversible<br>(E.C.2.1.2.1) | 500.5 | Glycine, serine and threonine metabolism is affected upon addition of reactive species to the metabolic network (34) resulting in addition of folate derivatives to folic acid pool. |
| 2. | methylenetetrahydrofolate<br>dehydrogenase (NADP)<br>(E.C. 1.5.1.5) | 1000 | NADPH and serine metabolic pathways are also linked to folate metabolism (35), and get affected in presence of ROS, thereby generating positive flux values. |
| 3. | 2-Hydroxybutyrate:NAD<sup>+</sup><br>oxidoreductase<br>(E.C. 1.1.1.27) | 105.8 | Association between butyrate and folic acid pathways has not been established experimentally yet and needs to be validated. |

Perturbations in gut microbiota have been linked to inflammatory bowel disease, as well as, the cancers of colon and stomach (8). Similarly, the alterations in gut bacterial metabolites production can also contribute to the initiation or treatment of these metabolic disorders. For instance, folate deficiency has often been associated with increased risk of colon cancer. On the contrary, supplementing folic acid in patients diagnosed with cancer, resulted in formation of aberrant crypt foci and initiation of cancer in colon polyps (29). The quantitative effects of lower folic acid concentrations on cancer cells have not been studied so far. Surprisingly, the experimental findings showed that lower concentrations of folic acid had cytotoxic effects on HCT116 colon cancer cells.

The AgNP treated bacterial cultures also showed increased production of folic acid, as a consequence of increased oxidative stress. The supernatants of the AgNP treated bacterial cultures with increased folic acid content reduced the viability of HCT116 colon cancer cells. The folic acid concentrations in these cultures were equivalent to the optimal-cytotoxic concentration (0.5 to 1 μM) of synthetic folic acid that resulted in cell death. This result indicates that oxidative stress caused increased production of folate to optimal-cytotoxic levels, which resulted in decreased cancer cell viability.

## Conclusion

In our study, AgNPs when used at lower concentrations, showed no major cytotoxic effects on the *E. durans*. Instead, AgNP-generated oxidative stress modulated the folic acid levels in the microbe. This finding was supported by genome scale metabolic modeling. Further, metabolic association between folate and SCFA metabolic pathways was observed. Other key players to this were identified as amino acids, energy metabolites, and nucleotides. Lastly, folic acid, at lower concentrations was found to exert cytotoxic effects on HCT116 colon cancer cell line, thus, highlighting the potential of critical folate concentration for CRC treatment.

## Experimental procedures

### Bacterial culture and growth conditions

*Enterococcus durans*, a facultative aerobe, procured from MTCC (MTCC No. 3031) was used as the model organism. The bacterial culture was grown in shake flasks containing MRS broth at 37° C, 180 rpm, in a shaker (Scigenics Orbitek). The total cell concentrations at different time points were measured through optical density (cell scatter) at 600 nm (JASCO V-630 Spectrophotometer), and comparison with a standard plot of OD vs. cell concentration was done.

### Characterization of silver nanoparticles (AgNPs)

Silver Nanoparticles (AgNPs) were obtained from Sigma Aldrich (catalogue no. 7440-22-4). Size distribution analysis was carried out using dynamic light scattering and zeta potential was measured using Horiba Scientific nanopartica nanoparticle analyzer (SZ-100). Scanning electron microscope was used to study NP-bacteria interaction.

### Treatment of bacterial cells with silver nanoparticles (AgNPs)

The AgNPs were dispersed in medium using water bath sonicator, till the nanoparticles were dispersed in the solution. AgNPs at 25-250 ppm concentrations was used. The medium was then inoculated with the appropriate volume of the subculture (inoculum), such that the OD value at the zeroth hour was 0.1.

### Quantifying intracellular ROS concentrations

Intracellular RS were measured by following the procedures from literature (31). The fluorescent dyes 3’-(p-amino-phenyl) fluorescein (Invitrogen, USA) and dihydroethidium (Sigma-Aldrich, India) were used to detect hydroxyl radical and superoxide radical, respectively. Hydroxyl and superoxide radical concentrations were determined from calibration curves using the standards hydrogen peroxide and potassium superoxide, respectively. The concentration of superoxide and hydroxyl radicals was reported in nanomoles per gram-cell weight.

### Sample preparation for folate estimation

For analyzing intracellular folate concentrations, bacterial culture was harvested every 3 hours and culture volume corresponding to 10 OD was used for sample preparation. The bacterial pellet obtained on centrifuging the required volume was suspended in 1ml of milli Q water. It was then sonicated (Q Sonica sonicator), at amplitude of 70%, for a process time of 4 minutes (pulse on and off time being 2 seconds). The sonicated sample was then placed in a water bath at 100°C and subjected to heat for 5 minutes so as to release any folate bound to the folate binding proteins (FBPs). The cell free extract was obtained by centrifuging the sample. The supernatant was collected, filtered and used further for folate estimation.

For quantifying extracellular folate, released by the bacterial cells into the growth medium, 1ml of culture was collected every 3 hours and the filtered supernatant was used for HPLC (38).

### HPLC analysis of folate

Folic acid (Himedia, catalogue no. CMS175) was used as standard in estimation of folate in samples (cell extracts and bacterial supernatant). HPLC analysis of samples was performed using UFLC Shimadzu HPLC setup. C18 Hypersil column (25cm*4.6mm, 5 micron spherical packing) was used as the analytical column. 15% HPLC grade Carbinol in 0.05M KH_2_PO_4_was used as the mobile phase. The mobile phase and samples were first filtered through 0.46 micron filters before use. The mobile phase was then sonicated in a bath sonicator for 10 minutes for degassing the solution. The flow rate was maintained at 0.4 mL/min. Excitation wavelength of 295nm was used to analyze the folic acid peak (38). Folic acid standards were used at different concentrations (0-125 μM), and the retention time for the analyte was determined from the chromatographs.

### Treatment of HCT 116 with bacterial supernatant

HCT 116 (colon cancer cell line) was obtained from Dr. Bert Vogelstein, John Hopkins University, Baltimore, USA. The tumor cells were grown in Dulbecco’s Modified Eagle Media (DMEM), with 5% serum. The cells were seeded and grown in 96 wells plate and were treated with bacterial supernatant (control and 25ppm AgNPs), and different concentrations of synthetic folic acid. MTT cell proliferation assay was then performed to quantify cell viability on exposure to the drug after 48 hours of treatment. MTT (3-(4, 5-dimethylthiazol-2-yl) - 2, 5-diphenyltetrazolium bromide) is a widely used colorimetric method to measure cellular metabolic activity. The principle behind this method is based on the ability of nicotinamide adenine dinucleotide phosphate (NADPH)-dependent cellular oxidoreductase enzymes to reduce the tetrazolium dye MTT to its insoluble formazan, which has a purple colour. This assay, therefore, measures cell viability in terms of reductive activity as enzymatic conversion of the tetrazolium compound to water insoluble formazan crystals by dehydrogenases occurring in the mitochondria of living cells.

#### Modeling gut bacteria – ROS interplay Constraint based model formulation

##### S-matrix and steady state assumption

Constraint-based metabolic model of the target organism comprises of metabolic reactions, metabolites participating in these reactions, and genes that encode the enzymes catalyzing these reactions. On a metabolic network map, metabolites form the nodes, and reactions forms the links. Mathematically, these metabolic networks are represented as stoichiometric matrix (S matrix) of size m x n, where the row and column represent the metabolites (m) and the reactions (n), respectively. The values in each column denote the stoichiometric coefficient for every metabolite participating in a particular reaction. The negative value indicates metabolite consumption, whereas, positive value is for metabolite production. Non-participation of a metabolite in a reaction is marked zero in the matrix.

The stoichiometry intends to impose constraints on flow of metabolites (in the form of flux values i.e. mass) through the network. Flux through all the metabolic reactions in the network is represented by a vector v. COBRA models are simulated under the steady-state condition, mathematically defined by the following equation:

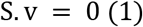

Where in, S is a sparse matrix because most biochemical reactions involve only a few different metabolites and any value of v that satisfies this equation falls in the null pace of S. The metabolic network is a system of mass balance equations at steady state (dx/dt=0), which signifies the total amount of a metabolite consumed is equal to the total amount of being produced in the system at steady state. Each reaction is also assigned lower and upper bounds, which define the range of allowable flux distributions of a system.

##### Flux Balance Analysis (FBA)

For a large scale metabolic model, generally, the number of reaction equations are greater than the number of metabolites in the system (n>m). Therefore, a whole solution space exists in such a scenario, there being no unique solution to this system of equations. In order to realistically narrow down this solution space, flux balance analysis (FBA) calculates and selects only those flux values that can together optimize some biologically relevant objective, like, finding metabolic reaction fluxes, simulation of growth on different substrates etc.

Mathematical formulation of FBA lies in optimizing (maximizing or minimizing) any objective function through linear programming (LP), which is represented by:

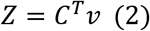

Here, c is the vector of weights that specifies to what extent each reaction contributes to the objective function. The output of FBA is given as a particular flux distribution, v, which either maximizes or minimizes the objective function Z. Since FBA doesn’t require kinetic parameters information, and can also be computed promptly for large networks, it can be utilized in studies involving characterization of various perturbations. However, it poses limitations in predicting metabolite concentration due to lack of kinetic data, and is suitable in determining fluxes at only steady state.

##### Flux Variability Analysis (FVA)

Flux Variability Analysis (FVA) is another tool to maximize or minimize the flux values for each reaction in the metabolic network, while simultaneously satisfying all imposed constraints on the optimized objective function value. It provides a span or range of allowable fluxes that can exist within the optimized solution space as defined by the linear program. Consequently, FVA provides useful insight into the network variability.

For our work, the unconstrained E. durans metabolic model was downloaded from Virtual Human Metabolism (VMH) database (https://vmh.uni.lu/) and then ROS reactions were added to the model.

rBioNet was used to add the different reactions (ROS reactions; missing transport and exchange reactions; sink and demand reactions) to the model. rBioNet enables the user to add these reactions in a quality controlled manner, by exempting sources of manual error (36). A total of 8 ROS reactions and 11 metabolites were added to the model. The resulting reconstruction (with ROS reactions) was then merged with downloaded metabolic model, thus formulating the ROS model (Supplement Table S1). Flux Balance analysis (FBA) was carried out on the newly added reactions individually to check if the reactions were blocked (carrying zero net flux). Blocked reactions might result from incomplete reaction information about the consumption of substrates or the generation of products. The blocked reactions were then resolved by adding complete metabolic reactions and pathways from literature). In case there were no evident supporting reactions pertaining to the blocked reaction, then the reaction was un-blocked (by adding demand or sink reactions for products and reactants respectively). In case of our *E. durans* model, seven demand reactions and one sink reaction was added. Upon un-blocking the newly added reactions, dead end metabolites were then identified. Dead end metabolites are the ones which are either produced or consumed but never both. These dead end metabolites were also resolved using literature information for adding the required reactions to the model. FVA was then performed that provides the flux values (minimum and maximum) through all the reactions in the metabolic network. Flux Span ratio (FSr) was calculated. FSr is the ratio between absolute net flux values of reactions for unconstrained model to absolute net flux values of reactions for constrained model with ROS reactions. The range for FSr is user defined. The reactions with FSr values in the range 0.8>FSr>2 were identified. These were the reactions that got affected due to the addition of ROS reactions to the model. MinNorm analysis was then carried out on the reactions of interest. MinNorm function in MATLAB is a tool for frequency estimation in a particular data vector, for a given function.

### Statistical analysis

All cultures and measurements were carried out in triplicates (each subjected to three technical replicates). Values have been reported as mean ± SD (please refer to individual results in the results section). One way ANOVA (level of significance =0.05) and Tuckey’s multiple comparison tests were carried out.

## Acknowledgements

This work was supported by the Department of Science and Technology, Government of India (grant no.: EMR/2016/002069) to RR, DK, and GKS. DST-INSPIRE Faculty award (DST/INSPIRE/04/2015/000036) to SS.

## Author contributions

PB, RR, DK, GKS, and SS conceived the various ideas and designed the experiments. PB performed all the experiments. All the authors wrote the manuscript.

## Supporting information

Supplement Table S1: Excel sheet containing details of the expanded *E.durans* model with ROS reactions.

## Footnotes

1 Abbreviations used are, CRC: colorectal cancer ROS: reaction oxygen species GEMs: genome-scale metabolic model COBRA: constraint-based reconstruction and analysis FBA: flux balance analysis FVA: flux variability analysis

## References

1. Sobhani, I., Amiot, A., Baleur, Y., Levy, M., Auriault, M. L., Van Nhieu, J. T., and Delchier, J. C. (2013) Microbial dysbiosis and colon carcinogenesis: Could colon cancer be considered a bacteria-related disease? Therap. Adv.Gastroenterol. 10.1177/1756283X12473674

2. Mármol I., Sánchez-de-Diego C., Dieste, P., Cerrada, E., and Yoldi, M. J. R. (2017) Colorectal carcinoma: A general overview and future perspectives in colorectal cancer. Int.J.Mol. Sci. 10.3390/ijms18010197

3. Dulal, S. and Keku, T. O. (2014) Gut microbiome and colorectal adenomas. Cancer J. 10.1097/PPO.0000000000000050

4. Gagnière, J., Raisch, J., Veziant, J., Barnich, N., Bonnet, R., Buc, E., Bringer, M. A., Pezet, D., and Bonnet, M. (2016) Gut microbiota imbalance and colorectal cancer. World J. Gastroenterol. 10.3748/wjg.v22.i2.501

5. Thursby, E. and Juge, N. (2017) Introduction to the human gut microbiota. Biochem. J. 10.1042/BCJ20160510

6. Guinane, C. M. and Cotter, P. D. (2013) Role of the gut microbiota in health and chronic gastrointestinal disease: Understanding a hidden metabolic organ. Therap. Adv. Gastroenterol. 10.1177/1756283X13482996

7. Friedland, R. P. and Chapman, M. R. (2017) The role of microbial amyloid in neurodegeneration. PLoSPathog. 10.1371/journal.ppat.1006654

8. Guarner, F. and Malagelada, J. R. (2003) Gut flora in health and disease. Lancet. 10.1016/S0140-6736(03)12489-0

9. Rodríguez, J. M., Murphy, K., Stanton, C., Ross, R. P., Kober, O. I., Juge, N., Avershina, E., Rudi, K., Narbad, A., Jenmalm, M. C., Marchesi, J. R., and Collado, M. C. (2015) The composition of the gut microbiota throughout life, with an emphasis on early life. Microb. Ecol. Health Dis. 10.3402/mehd.v26.26050.

10. Jandhyala, S. M., Talukdar, R., Subramanyam, C., Vuyyuru, H., Sasikala, M., and Reddy, D. N. (2015) Role of the normal gut microbiota. World J. Gastroenterol. 10.3748/wjg.v21.i29.8787

11. Keum, N. and Giovannucci, E. L. (2014) Folic acid fortification and colorectal cancer risk. Am. J. Prev. Med. 10.1016/j.amepre.2013.10.025

12. Ríos-Covián, D., Ruas-Madiedo, P., Margolles, A., Gueimonde, M., De los Reyes-Gavilán, C. G., and Salazar, N. (2016) Intestinal short chain fatty acids and their link with diet and human health. Front. Microbiol. 10.3389/fmicb.2016.00185

13. Hamer, H. M., Jonkers, D., Venema, K., Vanhoutvin, S., Troost, F. J., and Brummer, R. J. (2008) Review article: The role of butyrate on colonic function. Aliment. Pharmacol. Therap. 10.1111/j.1365-2036.2007.03562.x

14. Hinnebusch, B. F., Meng, S., Wu, J. T., Archer, S. Y. and Hodin, R. A. (2002) The effects of short-chain fatty acids on human colon cancer cell phenotype are associated with histone hyperacetylation. J. Nutr. 10.1038/nrc3610

15. Sak, K. (2012) Chemotherapy and Dietary Phytochemical Agents. Chemother. Res. Pract. 10.1155/2012/282570

16. Verhoef, M. J., Rose, M. S., White, M. and Balneaves, L. G. (2008) Declining conventional cancer treatment and using complementary and alternative medicine: a problem or a challenge? Curr. Oncol. 15(Suppl 2), s101–s106

17. Moorthi, C., Manavalan, R. and Kathiresan, K. (2011) Nanotherapeutics to overcome conventional cancer chemotherapy limitations. J. Pharm Sci. 10.18433/J30C7D

18. Zhang, T., Wang, L., Chen, Q. and Chen, C. (2014) Cytotoxic potential of silver nanoparticles. Yonsei Med. J. 10.3349/ymj.2014.55.2.283

19. Munger, M. A., Radwanski, P., Hadlock, G. C., Stoddard, G., Shaaban, A., Falconer, J., Grainger, D. W., and Deering-Rice, C. E.(2014) In vivo human time-exposure study of orally dosed commercial silver nanoparticles. Nanomedicine Nanotechnology, Biol. Med. 10.1016/j.nano.2013.06.010

20. Durán, N., Marcato, P. D., De Conti, R., Alves, O. L., Costa, F. T. M., and Brocchi, M. (2010) Potential Use of Silver Nanoparticles on Pathogenic Bacteria, their Toxicity and Possible Mechanisms of Action. J. Braz. Chem. Soc. 10.1590/s0103-50532010000600002

21. Lushchak, V. I. (2014) Free radicals, reactive oxygen species, oxidative stress and its classification. Chem.-Biol. Interact. 10.1016/j.cbi.2014.10.016

22. Devriese, L. A., Vancanneyt, M., Descheemaeker, P., Baele, M., Van Landuyt, H. W., Gordts, B., Butaye, P., Swings, J., and Haesebrouck, F. (2002) Differentiation and identification of *Enterococcus durans*, *E. hirae* and *E. Villorum*. J. Appl. Microbiol. 10.1046/j.1365-2672.2002.01586.x

23. Carasi, P., Racedo, S. M., Jacquot, C., Elie, A. M., Serradell, M. de losángeles, and Urdaci, M. C. (2017)Enterococcusdurans EP1 a promising anti-inflammatory probiotic able to stimulate sIgA and to increase *Faecalibacteriumprausnitzii* abundance. Front. Immunol. 10.3389/fimmu.2017.00088

24. Thiele, I. and Palsson, B. (2010) A protocol for generating a high-quality genome-scale metabolic reconstruction. Nat. Protoc. 10.1038/nprot.2009.203

25. Thiele, I., Heinken, A. and Fleming, R. M. T. (2013) A systems biology approach to studying the role of microbes in human health. Curr. Opin. Biotechnol. 10.1016/j.copbio.2012.10.001

26. Shoaie, S., Karlsson, F., Mardinoglu, A., Nookaew, I., Bordel, S., and Nielsen, J.(2013) Understanding the interactions between bacteria in the human gut through metabolic modeling. Sci. Rep. 10.1038/srep02532

27. Magnúsdóttir S., Heinken A., Kutt L., Ravcheev D. A., Bauer E., Noronha A., Greenhalgh K., Jäger C., Baginska J., Wilmes P., Fleming R. M. and Thiele I. (2017) Generation of genome-scale metabolic reconstructions for 773 members of the human gut microbiota. Nat. Biotechnol. 10.1038/nbt.3703

28. Quinteros, M. A., Cano Aristizábal, V., Dalmasso, P. R., Paraje, M. G. and Páez, P. L. (2016) Oxidative stress generation of silver nanoparticles in three bacterial genera and its relationship with the antimicrobial activity. Toxicol. Vitr. doi:10.1016/j.tiv.2016.08.007

29. Moody, M., Le, O., Rickert, M., Manuele, J., Chang, S., Robinson, G., Hajibandeh, J., Silvaroli, J., Keiserman, M. A., Bergman, C. J., and Kingsley, K. (2012) Folic acid supplementation increases survival and modulates high risk HPV-induced phenotypes in oral squamous cell carcinoma cells and correlates with p53 mRNA transcriptional down-regulation. Cancer Cell Int. 10.1186/1475-2867-12-10

30. Pieniz, S., Andreazza, R., Anghinoni, T., Camargo, F. and Brandelli, A. (2014) Probiotic potential, antimicrobial and antioxidant activities of *Enterococcus durans* strain LAB18s. Food Control, 10.1016/j.foodcont.2013.09.055

31. Menon, K. R., Balan, R. and Suraishkumar, G. K. (2013) Stress induced lipid production in *Chlorella vulgaris*: Relationship with specific intracellular reactive species levels. Biotechnol. Bioeng. 10.1002/bit.24835

32. Rao Y. M. and Sureshkumar G. K. (2001) Improvement in bioreactor productivities using free radicals: HOCl-induced overproduction of xanthan gum from *Xanthomonascampestris* and its mechanism. Biotechnol. Bioeng. 10.1002/1097-0290(20010105)72:1<62::AID-BIT9>3.0.CO;2-9

33. Ventura, M., O’Flaherty, S., Claesson, M. J., Turroni, F., Klaenhammer, T. R., van Sinderen, D., and O’Toole, P. W. (2009) Genome-scale analyses of health-promoting bacteria: Probiogenomics. Nat. Rev. Microbiol. 10.1038/nrmicro2047

34. Piper, M. D., Hong, S. P., Ball, G. E. and Dawes, I. W (2000) Regulation of the balance of one-carbon metabolism in *Saccharomyces cerevisiae*. J. Biol. Chem. 10.1074/jbc.M004248200

35. Fan, J., Ye, J., Kamphorst, J. J., Shlomi, T., Thompson, C. B., and Rabinowitz, J. D. (2014) Quantitative flux analysis reveals folate-dependent NADPH production. Nature. 10.1038/nature13236

36. Thorleifsson, S. G. and Thiele, I. (2011) rBioNet: A COBRA toolbox extension for reconstructing high-quality biochemical networks. Bioinformatics. 10.1093/bioinformatics/btr308

37. Fu, P. P., Xia, Q., Hwang, H. M., Ray, P. C., and Yu, H. (2014) Mechanisms of nanotoxicity: Generation of reactive oxygen species. J. Food Drug Anal. 22, 64–75

38. Jose, S., Bhalla, P., and Suraishkumar, G. K. (2018) Oxidative stress decreases the redox ratio and folate content in the gut microbe, Enterococcus durans (MTCC 3031). Sci Rep. 8, 12138

